# An evidence-based network approach to recommending targeted cancer therapies

**DOI:** 10.1101/605261

**Authors:** Jayaram Kancherla, Shruti Rao, Krithika Bhuvaneshwar, Rebecca B. Riggins, Robert A. Beckman, Subha Madhavan, Héctor Corrada Bravo, Simina M. Boca

## Abstract

In this work, we introduce CDGnet, an evidence-based network approach for recommending targeted cancer therapies, available as a user-friendly informatics tool. Our approach can be used to expand the range of options of targeted therapies for cancer patients who undergo molecular profiling. It considers biological pathway information specifically by looking at downstream targets of oncogenes and is personalized for individual patients via the user-inputted molecular alterations and cancer type. CDGnet integrates disparate sources of knowledge and provides results in a number of easily-accessible and usable forms, while separating targeted cancer therapies into categories in an evidence-based manner.

## Introduction

In today’s era of cancer precision medicine, therapeutic interventions are often tailored to an individual’s tumor molecular profile, in addition to traditional considerations including age, sex, cancer stage, medical and treatment history. The term “molecular profiling” is often used to refer to some test that considers one or more biomarkers. These biomarkers may be either genetic characteristics or mRNA or protein expression values. Genetic characteristics include point mutations, insertions, deletions, duplications, gene fusions and rearrangements. They may be either germline (inherited, present in normal tissue) or somatic (present in cancer cells but not normal tissue). Expression values refer to the expression of mRNA or protein in tumors, either in comparison to other tumors or to adjacent normal tissue. Typically, tumor molecular profiling is used when a patient has few or no standard treatment options left. However, for some tumor types, it is now routine to check for specific molecular features to decide on a targeted treatment plan. For example, KRAS-wild type colorectal cancer is generally treated with EGFR inhibitors^1^, ER-positive (ER+) breast cancer with aromatase inhibitors or antiestrogens such as tamoxifen or fulvestrant, and HER2-positive breast cancer with monoclonal antibodies trastuzumab and pertuzumab, tyrosine kinase inhibitors such as neratinib, or antibody-toxin conjugates such as trastuzumab-DM1.^2^ In many cases, if there is no FDA approved targeted therapy for a specific tumor type, clinicians may either recommend an off-label therapy that is prescribed for their alteration in another tumor type, or enrollment in precision medicine clinical trials (e.g. basket, umbrella, targeted therapy trials).

To make such decisions about off-label therapy recommendations, clinicians have to sift through vast amounts of literature and clinical databases to determine the clinical utility of variants identified through molecular profiling to decide on the appropriate treatment option for their patients. The same is true for clinical translational scientists considering relevant therapeutic approaches to evaluate in model systems or in humans, either utilizing single agents or combinations. In this setting, the number of possible molecular profiles that may be relevant and the number of experimental agents creates a combinatorial explosion of research possibilities among which prioritization is needed. Several efforts are ongoing to capture, standardize and share clinically relevant variants identified through molecular diagnostic tests between several public, academic, and private institutions,^3–5^ though challenges remain with synthesizing evidence in a manner that is both systematic and timely^6,7^. Our goal in this work is to expand the range of options of targeted therapies for cancer patients who undergo molecular profiling by developing CDGnet (Cancer-Drug-Gene network), a user-friendly, evidence-based approach that accounts for downstream effects within pathways in cancer and is personalized for the individual patient. Our tool, which uses the shiny framework with an R backend^8^, is available at: http://epiviz.cbcb.umd.edu/shiny/CDGnet/. We incorporate pathway information specifically by looking at downstream targets of oncogenes, which are genes that are constitutively activated in cancer^9^. This is explained in the cartoon in Figure 1: If an oncogene in a biological pathway is activated, targeting genes and proteins that are found upstream may no longer be effective, leading to a focus on downstream targets. This includes the scenario of EGFR inhibitors for KRAS-wild type colorectal tumors: The EGFR protein triggers a signaling cascade in cancer, which may be blocked by anti-EGFR drugs; however, this is only effective if KRAS, which is downstream of EGFR, is not mutated. This leads to a lack of effectiveness of therapies that block EGFR. As a result, colorectal cancer patients are typically tested for KRAS mutations and EGFR inhibitors are only prescribed to individuals without specific KRAS mutations in codons 12 and 13. A comprehensive characterization of untreated colorectal tumors estimated that 43% of non-hypermutated tumors had KRAS mutations and these mutations were generally oncogenic activating mutations^10^, which means that a large percent of colorectal cancer patients are left with few therapeutic options. Our framework and tool are seeking to remedy this issue.

**Figure 1:**
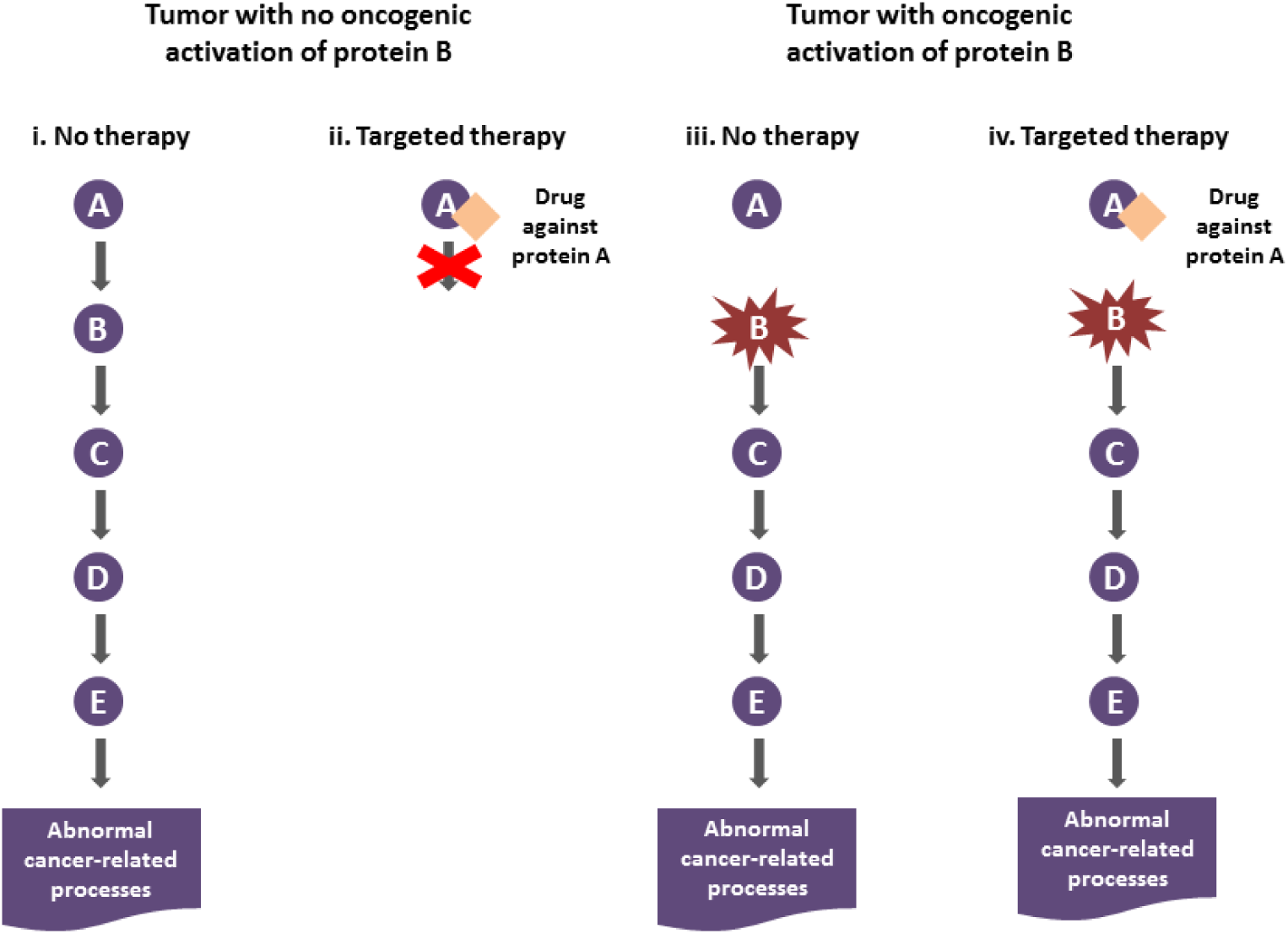
Simplified cartoon showing the reasoning behind looking at downstream targets of activated oncogenes. A simple pathway is shown that consists of 5 proteins, A, B, C, D, E, with A activating B, B activating C, etc, with the final activation of E leading to various abnormal cancer-related processes. Panels i and ii show the scenario where a tumor has no oncogenic activation of protein B, whereas panels iii and iv show the scenario where protein B has gained an oncogenic mutation that renders it constitutively active. If there is no oncogenic activation of protein B, then targeting protein A, as in panel ii, may be effective in stopping cancer growth. However, if there is oncogenic activation of protein B, this means that, in particular, it is not necessary for protein A to activate protein B, so that targeting protein A is not effective for turning off the pathway.

## Methods

### Overview of methods for generating patient-specific networks

The user inputs into CDGnet are the specific alterations found in a patient’s tumor and the patient’s cancer type. Part of the landing page is shown in **Figure 2**. These data are then integrated with: biological networks relevant to the cancer type (from the KEGG database^11^), FDA-approved targeted cancer therapies and indications (curated from DailyMed therapy labels^12^), additional gene-drug connections in the form of drug targets (from the DrugBank database^13^), information on whether a gene is an oncogene i.e. overactivated in cancer (from KEGG). Users may consider different data sources by using the code at https://github.com/SiminaB/CDGnet/ directly, for example by considering the oncogenes from a recent comprehensive characterization of The Cancer Genome Atlas (TCGA) projects^14^. Currently, the biological networks we consider are the cancer-specific pathways in KEGG and thus, for now we are also restricting the cancer types to those that have KEGG pathways. We have developed 4 different therapy categories that can be prioritized for a patient, given their specific tumor alterations, ordered from “most evidence that therapy works” to “least evidence that therapy works”:

**Figure 2:**
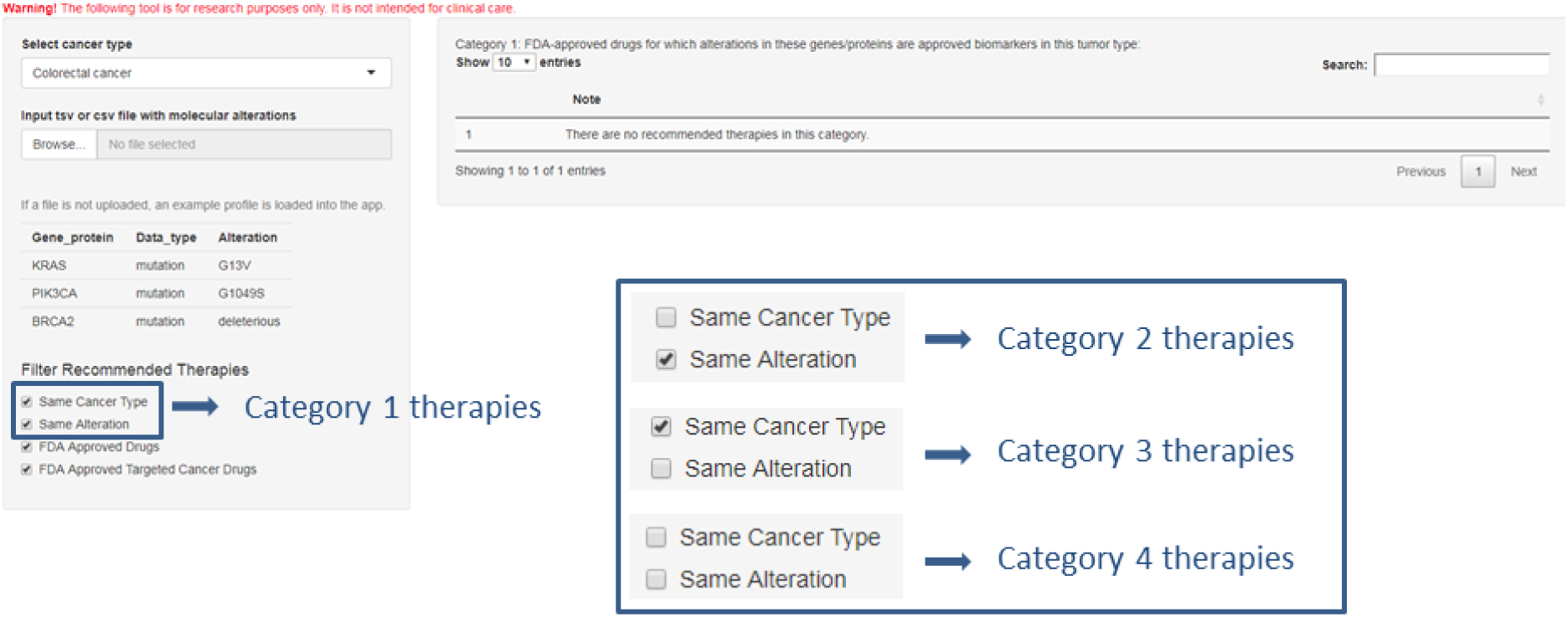
Part of the landing page, which shows how users can select the cancer type and either input a tab-separated or comma-separated file or use the example data. The inset shows how under “Filter Recommended Therapies,” combinations of the first 2 checkboxes lead to the 4 different categories of therapy recommendations described below. Removing one or both of the last 2 checkboxes expands the range of therapies in Categories 3 and 4 beyond FDA-approved drugs, respectively FDA-approved targeted cancer drugs.

1. FDA-approved drugs for which the input genes/proteins are biomarkers for their tumor type;
2. FDA-approved drugs for which the input genes/proteins are biomarkers in other tumor types;
3. Drugs which have as targets the input genes/proteins or as biomarkers/targets other genes/proteins that are downstream of the input oncogenes when considering the pathway corresponding to this tumor type; and
4. Drugs which have as biomarkers/targets other genes/proteins that are downstream of input oncogenes when considering the pathways corresponding to other tumor types.

In categories 3 and 4, users have the option to consider only FDA-approved targeted cancer therapies, all FDA-approved therapies, or all drugs in DrugBank; this allows for clinical researchers to consider increasing numbers of therapies only as needed, as opposed to being overwhelmed with a huge number of therapies from the start. For the clinician, this list reflects not only level of evidence but also practicality of obtaining the drug for use in their patient. Category 1 drugs will be readily available. Category 2 drugs will be readily available but may not be covered by the patient’s insurance, or may require considerable effort or justification to obtain coverage. Drugs that are not FDA approved may not be available unless their manufacturer has a compassionate use program, and then only with considerable effort.

We differentiate between targets and biomarkers because in many cases, due to complicated biological interactions, a therapy’s target may be different from the biomarker used to specify the indication, such as the case specified above with EGFR inhibitors being given for KRAS wild-type colorectal tumors, or CDK4/6 inhibitors being given for ER+ breast tumors. The general approach is presented in **Figure 3**. The options used on the landing page to obtain the different therapy categories are shown in **Figure 2.**

**Figure 3:**
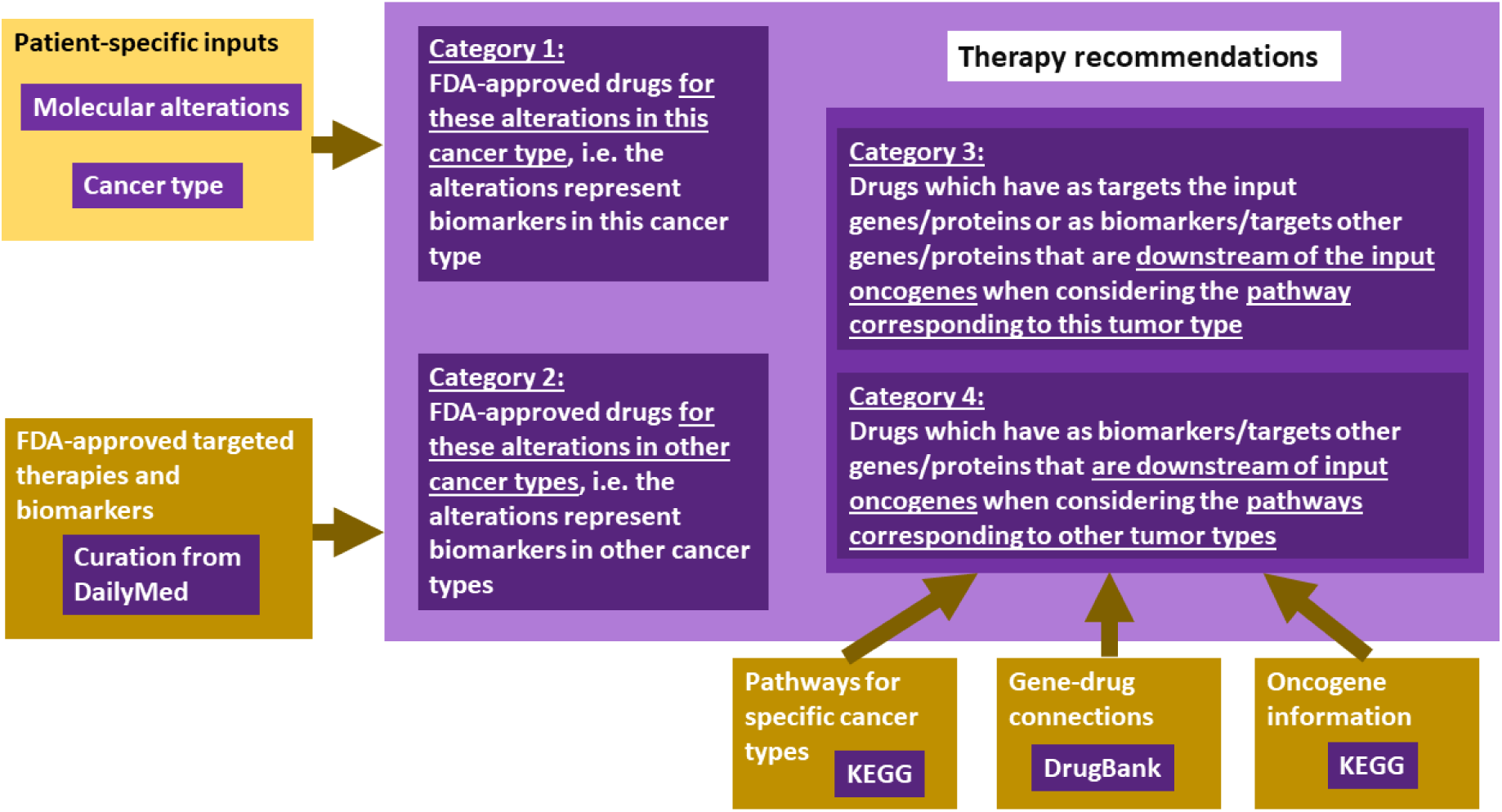
General approach for targeted therapy recommendations, including specific data sources.

Supplementary Table 1 gives the list of FDA-approved targeted cancer therapies and indications which was obtained by considering the targeted therapies listed by the NCI^15^ and looking up the corresponding labels at DailyMed^12^. In particular, the “Indications and usage” portion of the label was used to obtain the specific cancer type and the biomarker information, which is listed in the “Gene/Protein,” “Data type,” and “Alteration” columns; in the case of multiple biomarkers, these are listed on separate rows of the table. In cases where the biomarker indication is unclear, the lists of FDA companion diagnostic tests were also consulted^16,17^. Note that while some targeted therapies have specific biomarker indications, many do not. For example, ibrutinib (Imbruvica) is a targeted therapy, given for a number of subtypes of leukemia/lymphoma, but not for a specific indication. If there is no biomarker indication, this is noted as a “*” in the table under the “Gene/Protein” column. The therapies are then cross-referenced with DrugBank to obtain the targets for both the therapies with biomarker indications and those without indications. The biomarkers and targets obtained in these ways are checked against downstream targets from KEGG cancer-specific pathways – which were downloaded, parsed, and with identifiers converted using the KEGGREST,^18^ KEGGgraph,^19^ and org.Hs.eg.db^20^ Bioconductor packages respectively – and the information input by the user, with the gene/protein names being normalized via the rDGIdb package, which is a wrapper for the DGI database^21,22^.

In order to obtain the list of FDA-approved drugs, we used the data files from the official Drugs@FDA resource.^23^ Drugs@FDA contains several tab separated files (tsv) that include information on the submission, review and approval process for various drugs. We use the ‘products’ (list of all drugs) and ‘submission’ (review process for all drugs) files from these files to filter for only drugs that are “Approved” (AP) or “Tentatively Approved” (TA). The Drugs@FDA resource contains a list of all drugs approved since 1939 and some of these might be discontinued. As a result, we use the ‘marketingstatus’ file to remove any discontinued products from the list. The R scripts to parse and filter the Drugs@FDA data files are available in our GitHub repository.

### Shiny app and visualization

For each of the 4 categories detailed above, a sortable and searchable table of therapies is output with the FDA-approved indications; for categories 3 and 4 network visualizations are also shown. **Figure 4** shows a Sankey flow diagram representation which focuses on the flow of evidence between drug-gene and gene-gene connections, enabling an intuitive visualization from the molecular profile to the inferred targets and recommended therapies**. Figure 5** shows a portion of the sortable and searchable corresponding table. The path column represents the pathway between the altered gene/protein and the gene/protein that is a biomarker or target; the alteration column represents the biomarker for an FDA-approved indication, if this exists, in which case the tumor for which it is approved is also listed; the predicted effect column is “sensitive” if the alteration column is not empty, and “target” if the drug targets the protein according to the DrugBank data.

**Figure 4:**
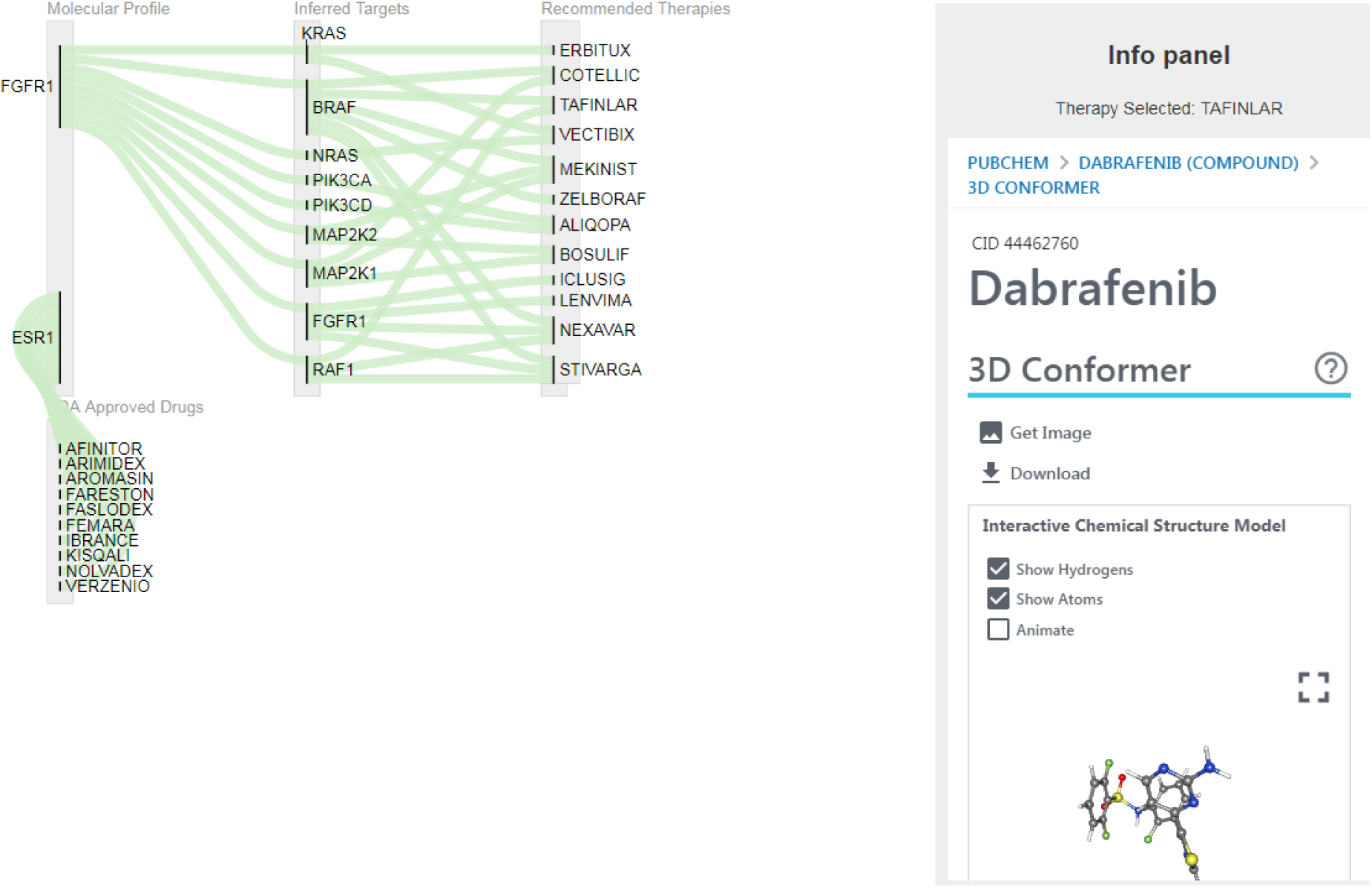
Sankey flow diagram, focusing on the flow of evidence between drug-gene and gene connections for a putative patient with ER+ breast cancer and FGFR1 overexpression, showing Category 3 recommendations, namely targets downstream of FGFR1. Therapies can be clicked to obtain a panel with PubChem information.

**Figure 5:**
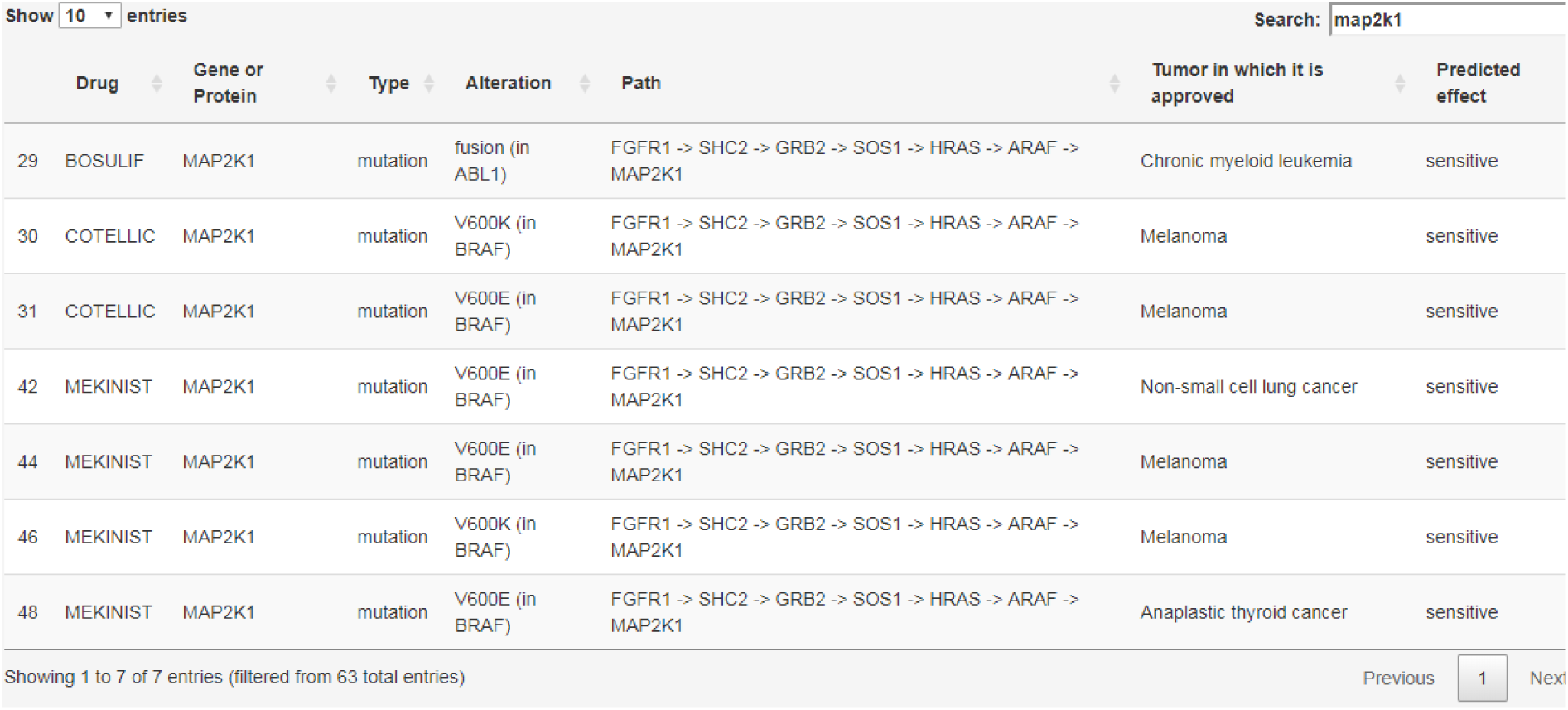
Part of the sortable, searchable table for therapies in Category 3 for a putative patient with ER+ breast cancer and FGFR1 overexpression, showing the subset of therapies that target MAP2K1.

An architecture diagram for our system is shown in **Figure 6**. We use shiny, an R package/framework for creating interactive and standalone web applications directly from R^8^. Shiny applications can run on a webpage or can be embedded in RMarkdown documents to build interactive dashboards. They use the same technology that powers web applications - HTML and JavaScript - and allow users to create intuitive and interactive user interfaces and prototypes with an R computational backend.

**Figure 6:**
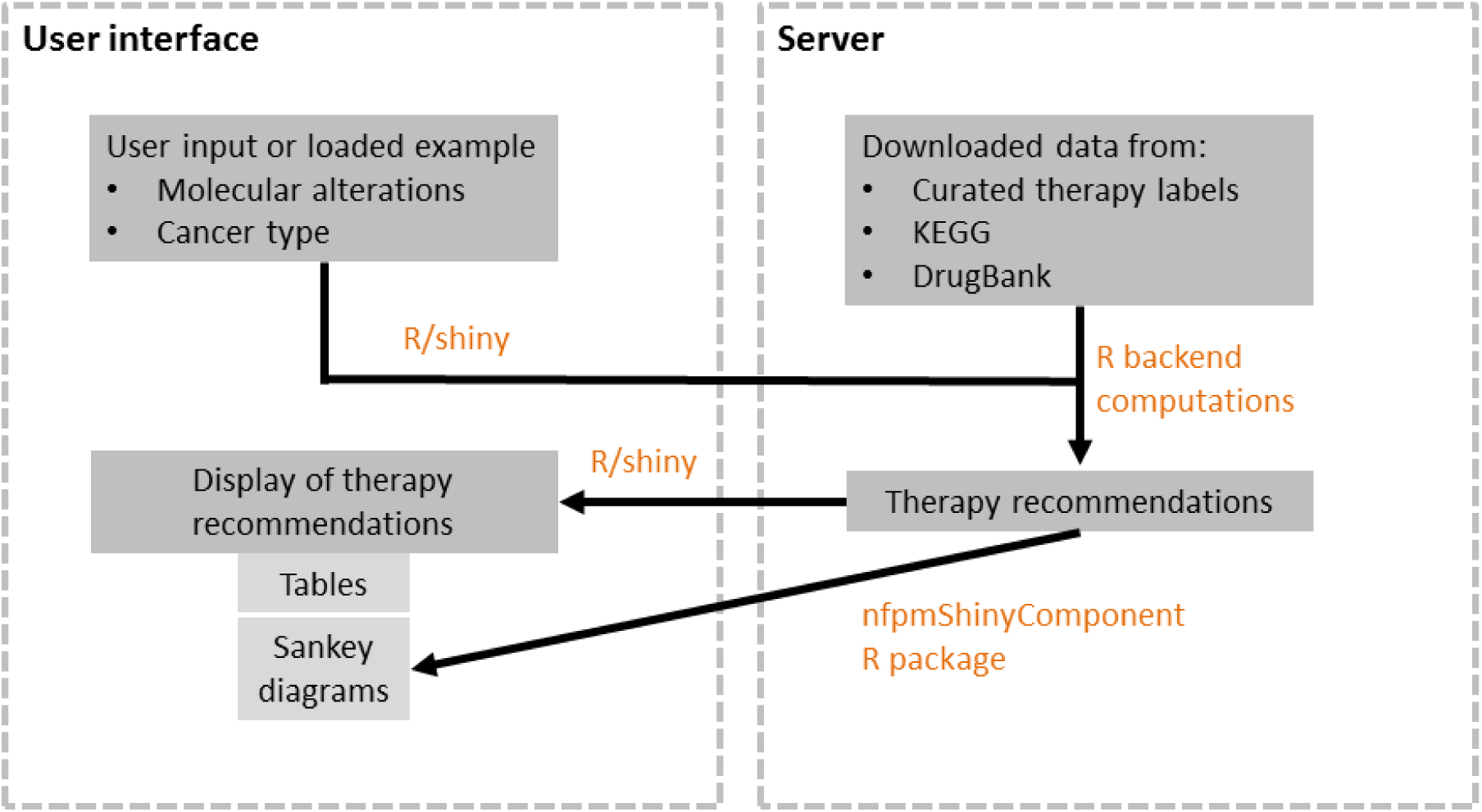
Architecture diagram for our system.

To support interactive Sankey charts inside shiny applications, we developed a shiny web component for visualizing Sankey flow diagrams, available to download as an R package at https://github.com/jkanche/nfpmShinyComponent. Web components are custom HTML elements that are natively extensible, reusable and can be integrated with any framework that supports HTML. The Sankey visualization uses a custom three column layout to organize nodes in the graph – 1) Molecular Profile and FDA approved drugs 2) Inferred Targets and 3) Recommended Therapies – and intuitively focuses the user on the flow of evidence from input parameters to recommended therapies. The Sankey visualization also contains an information panel that displays evidence related to a pathway connection or a drug when a user selects/clicks on an edge or node. Selecting an edge shows the downstream pathway information used for inference. Selecting a recommended therapy displays the structure of the drug and linked publications from PubChem,^24^ using PubChem widgets. The Sankey visualization is built on top of d3.js^25^, a data visualization library for JavaScript to build highly customizable and interactive visualizations.

## Results

We will now consider the scenario of a patient who has ER+ breast cancer. ER+ breast cancer, generally treated with aromatase inhibitors or antiestrogens, employs an array of mechanisms that permit escape from these therapies. These include amplification or upregulation of fibroblast growth factor receptor 1 (FGFR1), which is amplified in ∼13% of ER+ tumors from The Cancer Genome Atlas^26^–^28^ and leads to ligand-independent ER activation.^29^ FGFR activity has also recently been shown to confer resistance to CDK4/6 inhibitors in ER+ breast cancer.^30^ Pan-FGFR antagonists have been combined with endocrine therapies in prior clinical studies (e.g. CTKI258A2210), but the efficacy of this combination was minimal, even in patients pre-selected for alterations in the FGFR pathway.^31^ A potential underlying explanation for this lack of benefit is that FGFR alterations impinge upon downstream signaling networks shared by many other receptor tyrosine kinases. **Figure 4** shows CDGnet recommendations for a breast cancer patient with overexpression of both ESR1 (gene encoding ER) and FGFR1, when considering only FDA-approved targeted therapies. Therapy recommendations include PIK3CA, MAPK, and RAF inhibitors, which may have utility in this context, along with the standard targeted therapies prescribed for ER+ breast cancer. **Figure 5** shows the subset of the corresponding table that consists of FDA-approved MAP2K1 inhibitors, which are approved for either ABL1 fusions or specific BRAF mutations in chronic myeloid leukemia, respectively melanoma, non-small cell lung cancer, and anaplastic thyroid cancer.

## Discussion

We developed the CDGnet tool, using an approach that considers biological pathways and connections between genes, proteins, and drugs, to prioritize targeted therapies for cancer patients. Our approach integrates many disparate sources of knowledge and provides results in an easily-accessible and usable format. Using our tool, clinicians and clinical researchers are able to quickly obtain information on the FDA approved therapies (Category 1) and potential off-label therapies (Category 2) associated with a patient’s molecular profile. Our definition of Categories 1 and 2 in CDGnet are in alignment with the Tier I and II evidence level classifications recommended by the Association for Molecular Pathology (AMP), American College of Medical Genetics and Genomics (ACMG), American Society of Clinical Oncology (ASCO), and College of American Pathologists^32^. However, CDGnet’s Categories 3 and 4 are unique to our evidence-based network approach and enable clinicians and clinical researchers to evaluate additional targeted therapy options based on their cancer patient’s profile. It is important to note that the targeted therapy recommendations in Categories 3 and 4 have lower evidence levels and may or may not have proven clinical significance in ongoing clinical trials. However, by examining the downstream targets of candidate biomarkers, clinical researchers can derive key insights on potential biological pathways that can be targeted by different cancer therapies. Based on the level of evidence, the clinical actionability of these pathways can be further tested in a laboratory or clinical trial setting. Additionally, there is a growing field of research related to drug-target interactions and drug repositioning using network-based models,^33–^ 36 which may in the future be integrated with our tool.

We aim to further enhance the data that drives the CDGnet tool by incorporating relevant information from additional precision oncology efforts, tools and resources. Users who download or connect to these resources may currently use them in the context of our approach by modifying our code at https://github.com/SiminaB/CDGnet. Expert-curated precision oncology databases include Clinical Interpretations of Variants in Cancer (CIViC)^5^, Cancer Genome Interpreter^37^, OncoKB^38^, DEPO^39^, and PMKB^40^, while more general resources include ClinVar^41^. These additional sources may further strengthen the clinical annotations and evidence related to germline and somatic alterations in our database and provide options in between curated drug label information and DrugBank targets. CIViC (https://civicdb.org/) is an open access, open source, community-driven web resource that allows clinical interpretations of mutations related to cancer. Cancer Genome Interpreter (https://www.cancergenomeinterpreter.org/biomarkers) is an online tool that connects genes and drugs along with their effect and publication sources, not as a network format, but as a tabular format. OncoKB (https://oncokb.org/) is another online precision oncology knowledge base that contains information about the effects and treatment implications of specific cancer gene alterations. DEPO (dinglab.ddns.net/) contains druggable variant information such as drug therapy, evidence levels (FDA-approved, Clinical Trials, Case Reports, Preclinical), and the cancer types for intended treatments. The Precision Medicine Knowledgebase (PMKB) (https://pmkb.weill.cornell.edu/) provides information about clinical cancer variants and interpretations. We also note that we are using a simplified model for incorporating pathway information via the consideration of targets which are downstream of oncogenes – there are scenarios which we do not capture where upstream targeting can also be useful, for example in the case of a feedback loop.^42^–^44^ We will incorporate more complex information in further iterations of our tool.

Consortia such as the Clinical Genome (ClinGen) resource’s Somatic Cancer Working Group^3,45^ and the Global Alliance for Genomic in Health’s (GA4GH) Variant Interpretation for Cancer Consortium (VICC)^46^ have ongoing efforts to standardize and harmonize the expert-curated data in these different knowledge bases, with the goal to enhance the interoperability between these databases. We will align the future development of CDGnet with the guidelines and consensus frameworks developed by these consortiums. CDGnet can also serve as an informative tool for oncologists, molecular pathologists and genomic scientists who routinely participate in Molecular Tumor Board discussions.

Tools similar to CDGnet include PreMedKB^47^ and Drug Gene Interaction Network. PreMedKB (http://www.fudan-pgx.org/premedkb/) is an integrated precision medicine knowledgebase for interpreting relationships between diseases, genes, variants, and drugs. Drug Gene Interaction Network (http://seqome.com/drug-gene-network/) is a commercial tool offered by Seqome Inc that builds drug-gene interaction networks to predict clinical response from multi-omics datasets. The advantage of CDGnet over these tools is that our approach allows users to input specific alterations found in a patient’s tumor and cancer type, and outputs therapy options ordered based on priority. Such a personalized tool may eventually expand the range of options of targeted therapies for cancer patients in a clinical setting, a key goal of precision oncology.

## Supplementary Information

**Supplementary Table 1.** Curated FDA-targeted cancer therapies using label information, last updated on July 12, 2018.

## Acknowledgments

Funding for this work was provided through NIH grants R21CA220398 and supplement R21CA220398-02, U41HG009649, and P30CA051008 (via pilot award.)

